# Type I interferon drives a cellular state inert to TCR-stimulation and could impede effective T-cell differentiation in cancer

**DOI:** 10.1101/2024.03.28.587179

**Authors:** Dillon Corvino, Martin Batstone, Brett G.M Hughes, Tim Kempchen, Susanna S Ng, Nazhifah Salim, Franziska Schneppenheim, Denise Rommel, Ananthi Kumar, Sally Pearson, Jason Madore, Lambross T. Koufariotis, Lisa Maria Steinheuer, Dilan Pathirana, Kevin Thurley, Michael Hölzel, Nicholas Borcherding, Matthias Braun, Tobias Bald

## Abstract

Head and neck squamous cell carcinoma (HNSCC) arises from the mucosal epithelium of the oral cavity, pharynx, or larynx and is linked to exposure to classical carcinogens and human papillomavirus (HPV) infection. Due to molecular, immunological, and clinical disparities between HPV+ and HPV-HNSCC, they are recognized as distinct cancer types. While immune checkpoint inhibition (ICI) has demonstrated efficacy in recurrent/metastatic HNSCC, response variability persists irrespective of HPV status. To gain insights into the CD8+ T-cell landscape of HPV-HNSCC, we performed multimodal sequencing (RNA and TCR) of CD8+ tumor-infiltrating lymphocytes (TILs) from treatment-naïve HPV-HNSCC patients. Additionally, we subjected cells to *ex vivo* TCR-stimulation, facilitating the tracing of clonal transcriptomic responses. Our analysis revealed a subset of CD8+ TILs highly enriched for interferon-stimulated genes (ISG), which were found to be clonally related to a subset of granzyme K (GZMK)-expressing cells. Trajectory inference suggests ISG transition via GZMK cells towards terminal effector states. However, unlike GZMK cells, which rapidly an effector-like phenotype in response to TCR stimulation, ISG cells remain transcriptionally inert. Consequently, ISG cells may impede effective T-cell differentiation within the TME. Although, the functional consequences of ISG cells are poorly understood, we revealed that they possess receptors and ligands enabling cell-cell communication networks with key TME immunomodulators such as dendritic cells. Additionally, ISG cells were found to be a core feature across various tumor entities and were specifically enriched within tumor tissue. Thus, our findings illuminate the complexity of T-cell heterogeneity in HPV-HNSCC and reveal an overlooked population of IFN-stimulated CD8+ TILs. Further exploration of their functional significance may offer insights into therapeutic strategies for HPV-HNSCC and other cancer types.

## Background

Head and neck squamous cell carcinoma (HNSCC) encompass cancers originating from the mucosal epithelium of the oral cavity, pharynx, or larynx. HNSCC is closely associated with myriad environmental and lifestyle factors such as air pollutants, tobacco, and alcohol consumption (Johnson et al. 2020). In addition, viral co-infection with human papillomavirus (HPV) is observed in a subset of HNSCC (∼32%) patients (Ndiaye et al. 2014). Interestingly, HPV+ HNSCC is associated with more favourable prognosis especially in early stage disease (Fung et al. 2017; Lassen et al. 2009; Ang et al. 2010). The clinical benefit of HPV status is thought to derive from HPV-specific immune responses and the intrinsic immunogenicity of HPV (Nelson et al. 2017; Andersen et al. 2014).

Standard-of-care treatment options for HNSCC include surgical resection, radiotherapy, and chemotherapy (Johnson et al. 2020). However, immunotherapy-based treatment approaches such as immune checkpoint inhibition (ICI), have shown significant clinical benefit in the recurrent/metastatic setting (Vos et al. 2021). In fact, immune checkpoint inhibition has been approved for first-line treatment of patients with recurrent/metastatic (R/M) HNSCC (Burtness et al. 2019). Unfortunately, response to immunotherapy varies significantly. Variable responses may, in part, be attributed to the immunosuppressive tumor-microenvironment (TME) commonly observed in HNSCC (Johnson et al. 2020). While it is generally accepted that HPV+ HNSCC shows more robust anti-tumor immune responses compared to HPV− HNSCC, recent immunotherapy trials did not find an association between HPV status and response (Sacco et al. 2021; Ferris et al. 2016). Given, that CD8+ T-cells are recognized as key drivers of anti-tumoral responses, a better understanding of the CD8+ tumor-infiltrating lymphocyte (TIL) heterogeneity in HPV-patients is needed to improve the treatment for this subgroup of HNSCC.

Interferons (IFNs) are pleiotropic cytokines primarily produced by immune and stromal cells in response to pathogens or malignant transformation. Three types of IFNs have been described, which differ by the distinct receptors they bind and the subsequent signaling cascades induced. Type I IFNs (IFN-I) have well described roles in both anti-viral and anti-tumor responses. In particular, IFN-I can directly inhibit tumor growth by inhibiting proliferation and inducing apoptosis. In addition, IFN-I can act indirectly to induce anti-tumor immune responses, for example via the activation of dendritic cells, natural killer cells or neutrophils (Bald et al. 2014). Simultaneously, IFN-I can reduce the pro-tumorigenic functions of regulatory T-cells and myeloid-derived suppressor cells (Yu, Zhu, and Chen 2022). In fact, IFN-I signaling is considered as a “third signal” of activation and important for naïve T-cell priming, activation, proliferation, and memory differentiation (Curtsinger and Mescher 2010). Thus, IFN-I is regarded as a crucial cytokine in facilitating cancer immunosurveillance and boosting the efficacy of cancer immunotherapies (Yu, Zhu, and Chen 2022; Fuertes et al. 2011; Diamond et al. 2011; Ruotsalainen et al. 2021). However, we have previously shown via genetic ablation, that IFN-I signaling is dispensable for the expansion and function of adoptively transferred tumor-specific CD8+ T-cells (Ruotsalainen et al. 2021). In addition, several studies also provide evidence that IFN-I signaling, at least in the later stages of anti-tumor immune responses, can promote pro-tumor changes and ultimately immune escape (Zhou et al. 2020). For example, IFN-I signaling is linked to expression of immune checkpoints, IL-10, Nos2 and the development of a T-cell exhaustion phenotype (Ruotsalainen et al. 2021; Chen et al. 2022; Sumida et al. 2022). Therefore, the effect of IFN-I signaling in the functional outcomes of tumor-infiltrating T-cells is multifaceted and requires further investigation.

Single-cell RNA sequencing (scRNA-seq) of immune cell subsets in cancer patients has enabled the high-resolution mapping of cellular heterogeneity. This methodology has been applied to the analysis of human T-cells in response to cancer immunotherapies (Sade-Feldman et al. 2018). However, traditionally this approach only focuses on assessing the transcriptional state of *ex vivo* isolated cells. Thus, capturing a snapshot of cellular transcriptomic landscape within the TME. Therefore, we leveraged an *ex vivo* perturbation via a short-term T-cell receptor (TCR) stimulation. Coupled with scRNAseq and single-cell TCR sequencing, we were able to study the clonal dynamics and evaluate the responsive potential of CD8+ tumor-infiltrating lymphocyte (TIL) subsets.

Herein, we sequenced over 11,000 resting and stimulated CD8+ TILs isolated from treatment-naïve HPV-HNSCC patients. As such, we were able to define *ex vivo* cellular states and their stimulation outcomes. Importantly, we identified a population of T-cells rich in IFN-stimulated genes (ISG). These ISG cells were found to be associated with an IFN-I signature and were specifically enriched within the tumor tissue of various tumor entities. Furthermore, these cells were found to be clonally related to a population of cells highly expressing granzyme K (GzmK). However, unlike the GzmK subset, ISG-cells were transcriptionally inert to stimulation and thus possibly possess a unique role within the TME. This study sheds light on the existence of this overlooked population and begins to investigate their functionality.

## Results

### Single-cell RNA sequencing of CD8+ TILs from treatment-naive HNSCC patients identifies exhausted and effector populations

CD8+ T-cells are key drivers of anti-tumor responses. However, there is substantial heterogeneity in CD8+ T-cell phenotypes within TIL populations. As such, we sought to explore the diversity of CD8+ TILs in HPV-treatment-naïve non-R/M HNSCC patients. We isolated live CD45+CD3+CD4-CD8+ from 8 patients using flow cytometry-based cell sorting and subjected half of those cells to *ex vivo* CD3/28 TCR stimulation. After 5 hours of stimulation, we performed single-cell RNA and TCR sequencing to simultaneously identify CD8+ TIL phenotypes and clonotypes. We thereby were able to profile transcriptional changes in response to TCR-based stimulation (Figure 1A).

**Figure 1:**
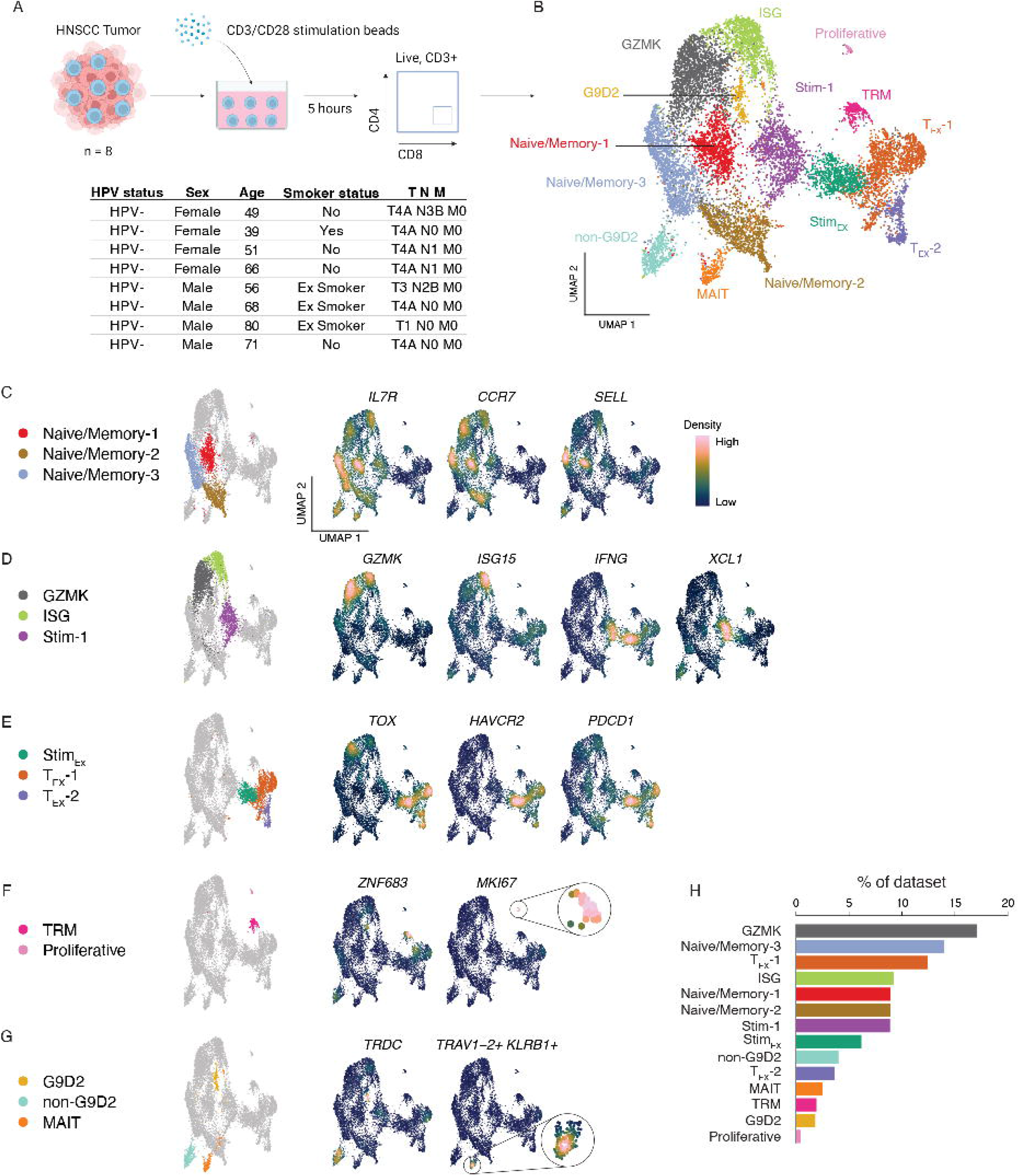
The transcriptional landscape of tumor-infiltrating CD8+ TILs in treatment-naive in head and neck squamous cell carcinoma (HNSCC) patients. (A) schematic detailing experimental setup used to generate the dataset. In brief, the tumors from eight head and neck squamous cell carcinoma (HNSCC) patients were digested and processed into a single-cell suspension. The cell suspension was cultured for 5 hours with or without CD3/CD28 T-cell stimulation. Subsequently, the cells were sorted for CD3+CD4-CD8+ T-cells and subjected to 10X single-cell sequencing. Key patient characteristics are listed in the table below the schematic. All patients were HPV negative, treatment naïve, and samples were from primary tumors. Schematic created with BioRender.com (B) UMAP projection of all cells that passed QC inclusion criteria. (C–G) UMAP projections highlighting (first column) clusters identified and subsequently the expression density of key genes used in their identification. (G) MAIT-cell identity is highlighted using the joint density expression of TRAV1-2 and KLRB1 (H) Barplot showing the frequency of each cluster identified as a proportion of the entire dataset.

Sequencing data from both unstimulated and TCR-stimulated samples were integrated and projected onto a unified UMAP space (Figure 1B). This resulted in 14 distinct clusters of CD8+ TILs with the majority of identified clusters evenly distributed across both unstimulated and stimulated conditions (Supplementary Figure 1A). Importantly, two new clusters emerged specifically post-TCR-stimulation (clusters Stimulated-1; Stim-1 and Stimulated-Exhausted; Stim_EX_). Three naïve/memory cell clusters were identified and annotated based on their expression of markers such as *IL7R*, *CCR7*, and *SELL* (Figure 1C). A cluster of cells expressing *GZMK* as well as *EOMES*, *NKG7*, *TNFRSF18* (encodes for GITR), and *CD69* was also identified (Figure 1D and data not shown). Additionally, a cluster of cells expressing high levels of various interferon-stimulated genes, including *ISG15*, *IFI6*, *IFIT3*, *MX1*, *ISG20*, *IFITM1*, *IFIT1*, *MX2*, and *OAS3* (Supplementary Figure 1B and data not shown) was recognized and annotated as the interferon-gene stimulated (ISG) cluster of cells (Figure 1D). The stimulated-1 (Stim-1) cluster from TCR-stimulated cells was enriched for the expression of immune effector molecules such as *IFNG*, *XCL1*, *XCL2*, *CRTAM*, *TNF*, *TNFSF14* (encodes for LIGHT) and *TNFRSF9* (encodes for 4-1BB) (Figure 1D and Supplementary figure 1B). Three exhausted cell clusters were also identified, all expressing high levels of canonical exhaustion markers such as, *TOX*, *HAVCR2*, *PDCD1* (encodes for Tim-3 and PD-1, respectively), *CTLA4*, *ENTPD1* (encodes CD39), and *TIGIT* (Figure 1E and Supplementary Figure 1B). One of these exhausted clusters was exclusively found post-TCR-stimulation and as such was designated as the Stimulated-Exhausted (Stim_EX_) cluster. A small cluster of tissue-resident memory (TRM) cells was identified based on the expression of canonical TRM markers such as *ZNF683* (encodes for HOBIT), *PRDM1* (encodes for BLIMP1), *ITGA1* (encodes for CD49A), *ITGAE* (encodes for CD103), and *CXCR6* (Figure 1F and supplementary figure 1B). A small population of proliferating cells was also identified by their enrichment for proliferation and cell cycle genes, notably *MKI67* (encodes for Ki-67) (Figure 1F).

### HNSCC TME is populated with unconventional CD8+ T-cells

We also identified three clusters of unconventional T-cells (Figure 1G and Supplementary Figure 1C). Two of these had gene expression patterns indicative of gamma delta (LJδ) T-cell subsets. The third cluster expressed markers corresponding with a mucosal-associated invariant T (MAIT) cell population. LJδ T-cell clusters could be differentiated based on the expression of TCR genes (Supplementary Figure 1D), marking the two clusters as the VLJ9Vδ2 T-cells (G9D2) and non-G9D2 populations. All unconventional T-cell populations expressed high levels of *CD3* and *CD8* as previously described (Kalyan and Kabelitz 2013; Gherardin et al. 2018) (Supplementary Figure 1E). Differential gene expression revealed that the G9D2 population expressed cytotoxicity markers such as *GZMA, GZMB, GZMH, GNLY, PRF1,* and *NKG7* (Supplementary Figure 1B and 1F). Non-G9D2 γδ T-cells expressed markers such as *TCF7*, *CD27*, *KLRD1*, and *SELL*. Analysis of differentially expressed transcription factors revealed that these three cell clusters had distinct and unique transcriptional regulatory programs (Supplementary Figure 1G). For example, G9D2 cells revealed specific enrichment for transcription factors *EOMES*, *ZEB2*, and *ZNF683* (encodes for HOBIT), while non-G9D2 cells were enriched for *ID3*, *IKZF2*, *TCF7*, and *BACH2*. Meanwhile, MAIT-cells demonstrated a distinct pattern of enrichment for transcription factors associated with the MAIT lineage, such as *RORA*, and *ZBTB16* (encodes for PLZF). Altogether, the unconventional T-cells, TRMs, and proliferative cells, cumulatively represented about ∼10% of TILs within the dataset (Figure 1H).

### Ex vivo TCR stimulation leads to the emergence of two transcriptionally distinct T-cell clusters

For further analysis, we removed the three unconventional T-cell clusters from the dataset and recalculated the UMAP coordinates (Figure 2A). We next sought to investigate the two cell clusters which predominantly arose from TCR-based stimulation. Importantly, both stimulation-induced clusters shared expression of a number of genes expected following TCR activation, including critical effector molecules such as *IFNG, GZMB* or *FASLG*, as well as activation markers as *ICOS* and *TNFRSF9* (encodes for 4-1BB) (Figure 2B). However, despite an overlap of activation-induced transcription, both stimulation-induced clusters showed distinct patterns of gene expression reminiscent of their origin (Figure 2C). For example, the Stim-1 cluster was enriched for genes such as *IL7R*, *XCL1*, *CD69*, *TNFSF14* (encodes for LIGHT), *CD28*, and *LTB*, whereas the Stim_EX_ cluster expressed high levels of exhaustion markers such as *TOX*, *LAG3*, *HAVCR2* (encodes for TIM-3) and *CD96*. These basal gene expression profiles seem to overlap with gene expression of other clusters of the dataset. For example, genes enriched in Stim-1 cluster were also highly abundant in Naïve/memory, GZMK, and ISG clusters, while genes expressed within the Stim_EX_ cluster were found enriched within the remaining two TEX clusters and to a lesser extend within the TRM and proliferating cell clusters. This overlap suggested the two stimulation-induced clusters may have arisen from different transcriptional states. To test this hypothesis, we used the single-cell TCR sequencing data to trace clonal populations between unstimulated and stimulated datasets.

**Figure 2:**
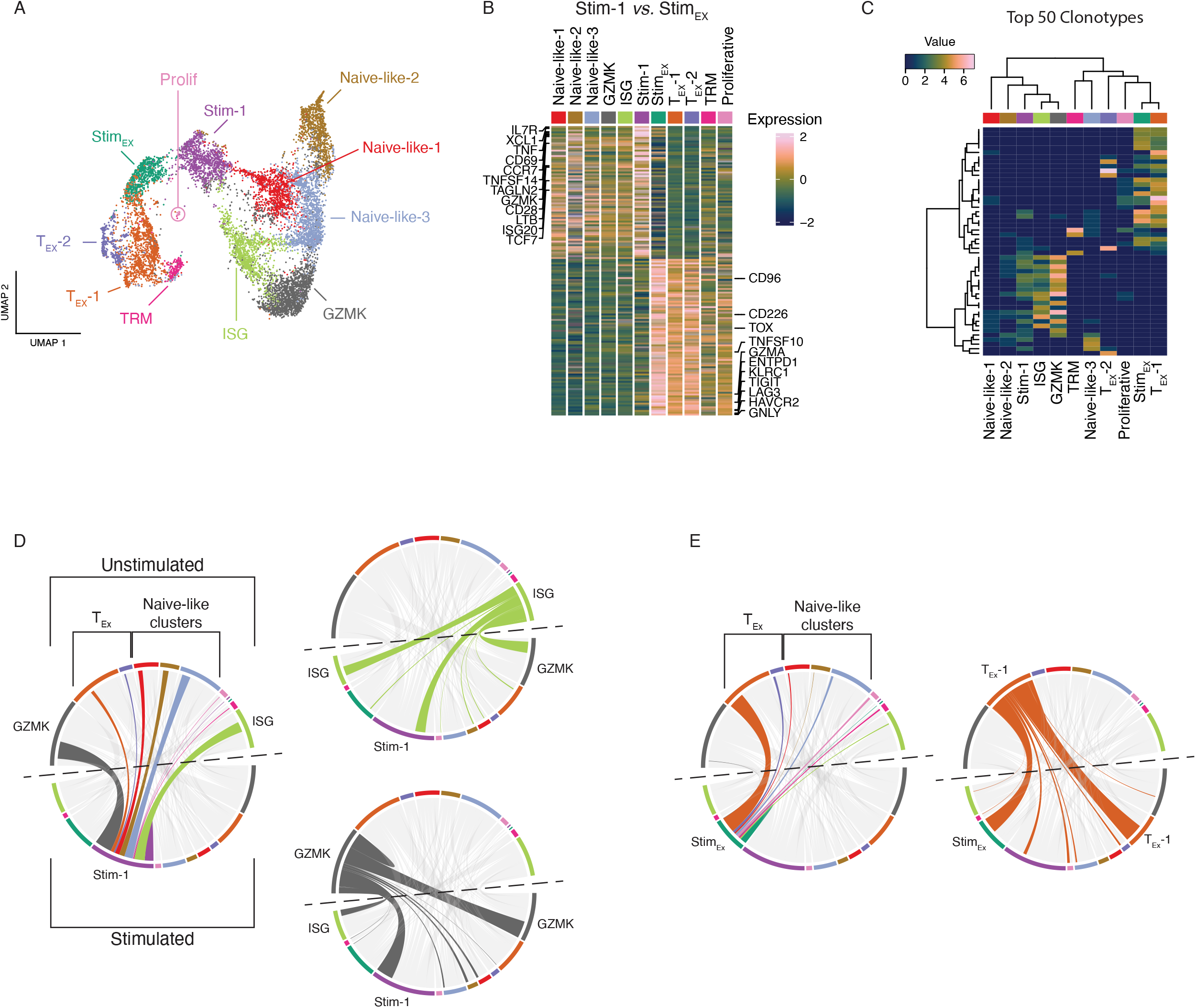
*Ex vivo* TCR stimulation induced transcriptional states develop from distinct unstimulated origins. (A) UMAP projection of CD8+ TILs identified in HNSCC patients after removal of unconventional T-cell subsets. (B) Heatmap of DEGs found to be upregulated (> 0.5 log2FC) in both stimulated-1 and stim-exhausted clusters, selected genes are annotated. (C) Heatmap of genes found to be significantly differentially expressed (>0.5 log2FC) between stim-1 and stim-exhausted clusters, selected genes are annotated. (D) Heatmap of the top 50 most abundant clonotypes found in CD8+ HNSCC TILs (ward.D2 clustering and binary distance function). (E) Stacked barplot showing the frequency of each clone size definition that is only found in the unstimulated sample (Unique to Unstimulated) or was also recovered post-stimulation (shared). Single (x = 1), small (1 < x < =5), medium (5 < x <=10), large (10 < x <=20) and hyperexpanded (20 < x <= 150). Where x = number of cells with exact CDR3 amino acid sequence. (F) Circos plots depicting the clonal overlap between clusters pre-(unstimulated; top arc) and post-stimulation (stimulated; bottom arc). Ribbons are coloured based on their unstimulated origin. Left column shows ribbons which connect to Stim-1 cluster whereas right column highlight ribbons that originate from ISG (top) or GZMK (bottom) clusters. (G) Same as (F) with left plot highlighted to show ribbons connecting with Stim-exhausted (Stim_Ex_) and ribbons in right plot highlighting those that originate from unstimulated T_Ex_-1 cluster.

An evaluation of the top 50 clonotypes observed in the dataset revealed an overlap between the Stim-1 and the ISG and GZMK clusters (Figure 2D). In contrast, the Stim_EX_ cluster shared many highly abundant clones with the T_EX_-1 cluster, indicating clonal overlap between these populations. To explore this further, we next traced clones pre- or post-stimulation to investigate the clonal overlap with respect to stimulation and cluster identity. However, this analysis relied on the assumption that clones were sufficiently represented in both pre- and post-stimulation datasets. Indeed, it was observed that when clones are represented in 2 or more T-cells (clone size small), >60% of clones are captured within the stimulated dataset (i.e shared) (Figure 2E). Therefore, we proceeded with tracing the transcriptional responses of shared T-cell clones by linking their cluster identity pre- and post-stimulation. We observed that cells from the Stim-1 cluster largely overlapped with unstimulated ISG and GZMK clusters (Figure 2F). Tracing unstimulated ISG clones, we observed clonal overlap that suggested stimulated ISG cells, either maintain their identity or adopt a GZMK or Stim-1 transcriptional phenotype. Similarly, unstimulated GZMK cells either retained GZMK identity or adopted ISG or Stim-1 transcriptional profiles post-stimulation. In contrast, clones from the Stim_EX_ cluster were predominantly found to overlap with unstimulated T_EX_-1 cluster with a minimal contribution from other unstimulated clusters (Figure 2G). As predicted, unstimulated T_EX_-1 cluster clones overlapped with stimulated Stim_EX_ or T_EX_-1 clusters. Interestingly, this analysis also revealed that TCR-stimulation was capable of inducing a gene signature associated with T-cell activation in a subset of transcriptionally terminally exhausted T-cells (TCF7-TOX+PD1+) (Figure 1E & Figure 2B, C)

### ISG cells largely retain their transcriptional identity upon TCR stimulation

To further understand the responsiveness to TCR stimulation across the dominant effector-like clusters, we isolated and projected them onto their own UMAP coordinates (Figure 3A). Subsequently, clones shared across pre- and post-stimulation datasets but whose cells were entirely contained within the ISG or GZMK clusters within the unstimulated dataset were identified (Figure 3B). This resulted in 26 and 53 unique clonotypes within unstimulated ISG or unstimulated GZMK clusters, respectively. Following TCR-stimulation, the majority of ISG T-cells retained their transcriptional identity (Figure 3C). In contrast, over 50% of unstimulated GZMK T-cells adopted a Stim-1 transcriptional identity following stimulation (Figure 3D), while the remaining proportion retained their GZMK identity. Interestingly, there was minimal adoption of an ISG signature following stimulation of GZMK clones. These data were further supported by pseudotime trajectory inference analysis, which revealed a trajectory of differentiation originating within the ISG cluster and transiting via GZMK population through to the Stim-1 cluster (Figure 3E). This trajectory was revealed using both a tree-based method (Slingshot) and a linear inference method (SCORPIUS; data not shown). Taken together, this data suggests a trajectory of ISG > GZMK > Stim-1, however, the transition from ISG > GZMK appears limiting as ISG cells were poorly responsive to TCR stimulation.

**Figure 3:**
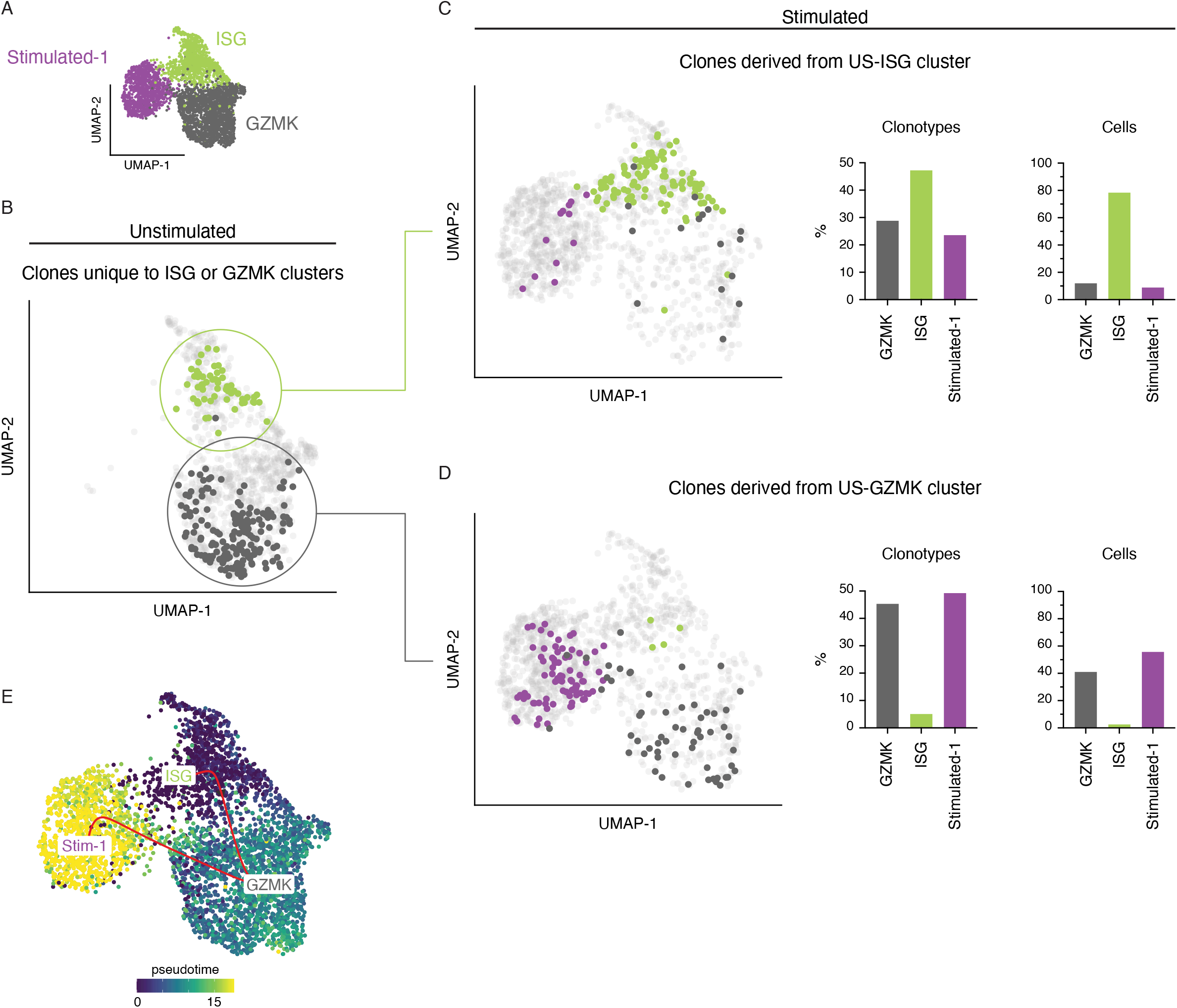
ISG cells are poorly transcriptionally responsive to TCR stimulation. (A) UMAP projection of Stimulated-1, ISG, and GZMK clusters both from unstimulated and stimulated datasets. (B) UMAP projection highlighting TCR clones uniquely found within unstimulated ISG cluster (green) or unstimulated GZMK cluster (black). (C) UMAP projection and quantification highlighting the distribution of unique US-ISG clones post-stimulation. Barplots quantify the frequency of cells post-stimulation. (D) same as (C) but for US-GZMK clones post-stimulation. (E) Pseudotime trajectory inference calculated using Slingshot, demonstrating potential progression of cells from an ISG state via GZMK through to Stim-1 phenotype.

### A type I interferon signature is associated with reduced transcriptional activity in ISG TILs

Given the diverse role of interferon signaling for the function of tumor-infiltrating T-cells, the relevance of ISG cells during tumor progression and immunotherapy remains elusive. We performed differential gene expression analysis and revealed a dominant signature enriched within the ISG population (Figure 4A). The top 10 differentially expressed genes identified within the ISG cluster were almost all found downstream of interferon signalling (Figure 4B). To understand the type of interferon signalling responsible, clusters were scored for genes contributing to a type I or type II interferon response (Figure 4C). Results showed the ISG cluster had enrichment for a type I, but not a type II interferon gene signature. Gene Ontology (GO) analysis was performed on the differentially up- or down-regulated genes within the ISG cluster relative to other clusters to unravel dominant biological processes associated with ISG cells. This analysis revealed a broad increase in translation related terms and type I IFN signalling responses (Figure 4D). Interestingly, down-regulated genes were enriched for GO terms associated with transcriptional regulation. This finding could explain our previous observation, that ISG cells poorly adopt new transcriptional states following TCR stimulation.

**Figure 4:**
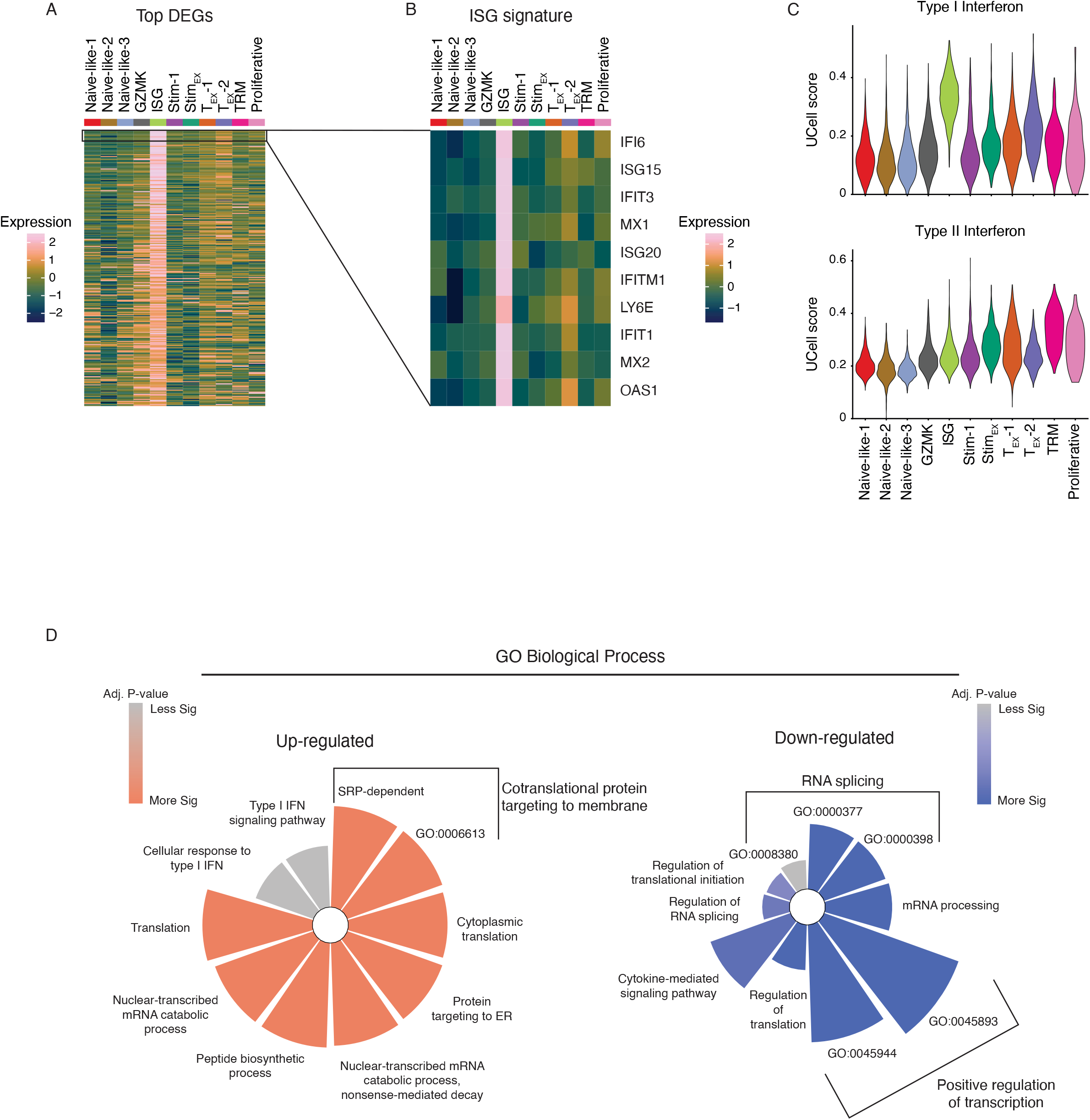
ISG cells are enriched for a type I interferon signature and are associated with reduced transcriptional. (A) Heatmap showing the Top up-regulated DEGs (> 0.25 Log2FC) identified in ISG cluster. (B) Heatmap showing top 10 DEGs identified in ISG cluster. (C) Violin plots of UCell scores for a type I interferon (top) or a type II interferon (bottom) gene signatures. (D) Gene ontology analysis for the top Up-regulated (left) and down-regulated (right) biological processes identified in the ISG cluster.

### ISG cells are enriched in CD8+ TILs across various tumor types

To establish whether ISG cells could be identified in other microenvironments, we generated a specific gene signature using the top 10 differentially expressed genes from ISG cells within our data set (Figure 4B). We next examined if this signature could identify ISG cells in a publicly available HNSCC dataset in which an ISG cluster had previously been reported (Cillo et al. 2020). Indeed, using our curated ISG-signature, we were able to correctly identify a cluster of cells enriched for type I interferon genes (Supplementary Figure 2A).

To better understand the abundance of ISG cells within CD8+ T-cells in healthy and malignant tissues, we scored cells from a pan-cancer dataset for our ISG signature (Nicholas Borcherding 2022). Indeed, we could identify a fraction of T-cells highly enriched for our ISG-signature (Figure 5A). Next, we assessed the frequencies of ISG cells across normal and tumor tissues. Here, we found ISG cells to be significantly increased in tumor tissues, relative to normal tissue (Figure 5B). ISG cells were most frequent in Ovarian and Esophageal tumor types but also detected to various degrees among all other tumor types assessed (Figure 5C). As expected ISG cells were solely enriched for type I but not type II IFN genes (Supplementary Figure 2B). We also assessed a COVID-19 dataset including some Influenza samples to determine if ISG cells are also enriched in the blood of virally infected patients (D. Wang et al. 2022). Indeed, in both conditions we observed a population of CD8+ T-cells enriched for our ISG signature (Supplementary Figure 2C) with a higher frequency in disease compared to healthy control samples (Supplementary Figure 2D), suggesting that the ISG cluster phenotype is not restricted to tumor immunity.

**Figure 5:**
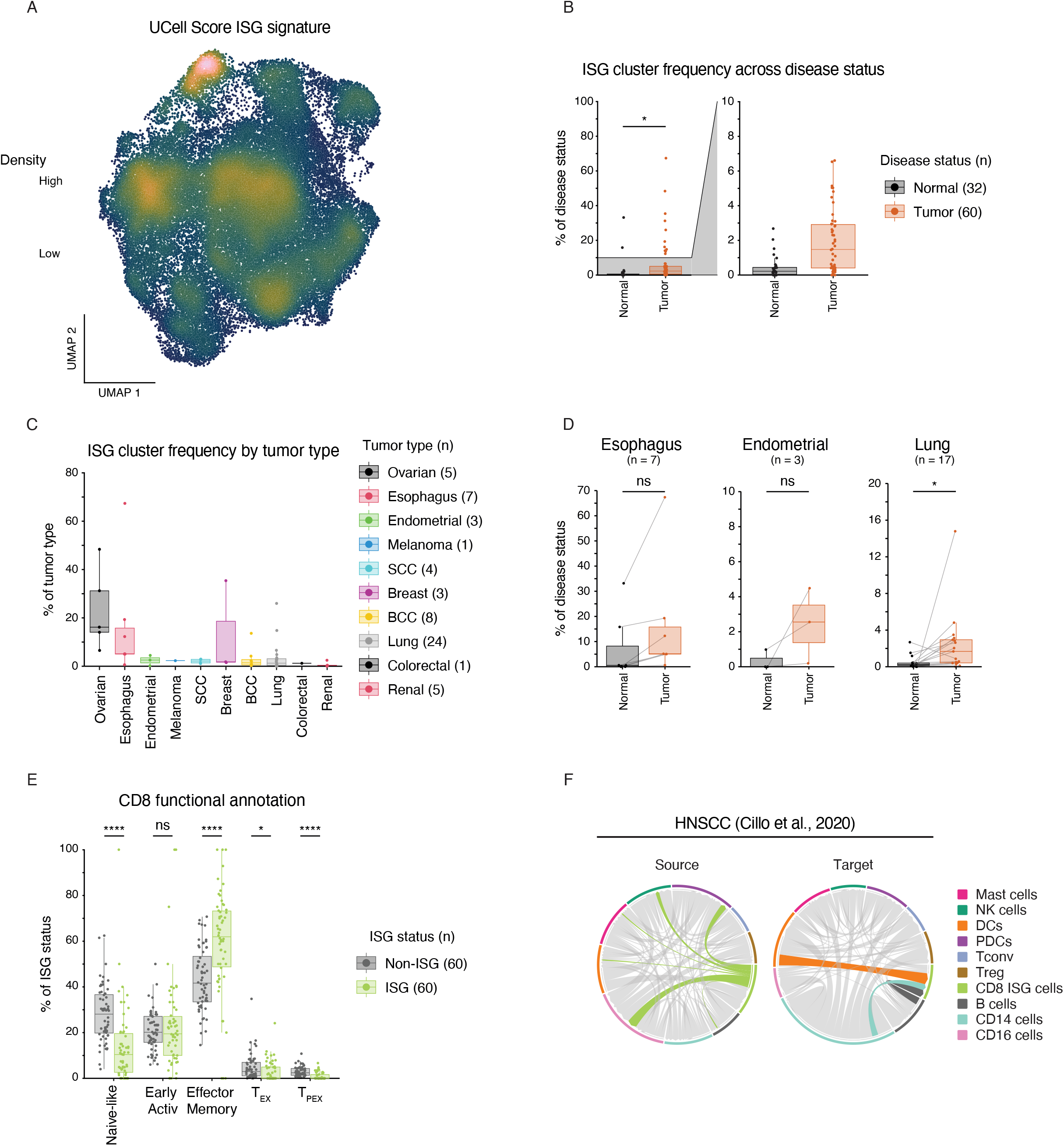
Cells with a type I interferon signature can be found across various tumor entities and are enriched within tumor tissue. (A) UMAP coordinates of CD8+ T-cells in a pan-cancer dataset overlaid with UCell score for ISG signature. (B) Boxplot showing ISG cluster frequency per donor across normal and tumor tissue samples. (C) Boxplot showing ISG cluster frequency within tumor samples per donor across tumor types within dataset. (D) Circos plots generated using the top 20 interactions for each source (left) or target (right) with ribbons highlighting interactions originating from ISG cluster (left) or terminating in ISG cluster (right), ribbons coloured by source. *p* value calculated using a two-tailed t-test. (n) value indicates the number of unique donors. ns = p > 0.05, * = p < 0.05, ** = p < 0.01, *** = p < 0.001, **** p < 0.0001.

Finally, to better understand the functional role of ISG cells within the TME, we employed cell-cell communication analysis. Utilising a published HNSCC dataset containing an array of immune cell subsets (Cillo et al. 2020), we revealed that ISG cells served as the source for interactions with CD16 positive cells, as well as with NK cells and plasmacytoid dendritic cells (PDCs) (Figure 5D). ISG cells were also found to be a target for DC, B cell, and CD14 cell interactions. Hence, this data suggests ISG cells interact with key innate immune cell subsets within the TME and thus potentially are important orchestrator of anti-tumor immunity.

## Discussion

HNSCC is a prevalent and complex disease with numerous etiological influences. For example, viral co-infection with HPV in Oropharyngeal HNSCC is associated with a better prognosis especially in early stage disease. As such, HPV-HNSCC presents as a more therapeutically challenging entity. Therefore, we sought to expand the knowledge base of CD8+ TIL landscape, specifically in treatment-naïve HPV-HNSCC patients. We employed a multimodal sequencing approach, together with an *ex vivo* TCR-stimulation, to facilitate tracing of transcriptional profiles and response capacity in CD8+ T-cell subsets.

Single-cell RNAseq of immune cell subsets has enabled in-depth mapping of the cellular heterogeneity of various disease conditions. However, traditionally this methodology only assesses the transcriptional state of cells *ex vivo*. Thus, capturing a snapshot of cellular transcriptomic landscape. Although, by leveraging an *ex vivo* perturbation coupled to sequencing approaches, others have ascertained both *ex vivo* profiles and their subsequent activation potentials. For example, a study by Szabo et al., 2019 performed *ex vivo* TCR-stimulation on T-cells isolated from several healthy donor tissues. The authors were able to define both conserved tissue signatures as well as the activation states of T-cells (Szabo et al. 2019). Using a similar approach, we included TCR sequencing to facilitate tracing of transcriptional responses within clonal populations of tumor-infiltrating T-cells. Notably, we observed two unique T-cell clusters specifically induced by TCR-stimulation. Transcriptional signatures and clonal overlap suggest these populations arose via stimulation of distinct *ex vivo* subsets. Importantly, we observed cells that displayed a transcriptional program of terminal exhaustion (*TCF7*-*TOX*+*PDCD1*+*TIM3*+), which retained substantial capacity to respond to TCR stimulation (Blank et al. 2019). These data posit transcriptionally exhausted cells may retain substantive capacity to respond to stimulation. Indeed, numerous scRNAseq studies have identified clusters of exhausted cells that simultaneously express high levels of effector molecules (Andreatta et al. 2021; Quah et al. 2023). These observations highlight the need for multimodal data approaches to identify prototypic exhausted T-cells while urging caution against defining exhaustion solely based on transcriptional profiles.

IFN-I signaling in CD8+ T-cells is associated with both anti- and pro-tumoral function (Zhou et al. 2020). Therefore, the clinical implications of an ISG-rich population is poorly understood. Substantial challenges impede the experimental investigation of these cells and as such our multi-modal sequencing approach has provided a comprehensive investigation of this population. Our analysis has revealed that CD8+ ISG cells are a common feature of solid malignancies and are specifically enriched within tumor tissue. Furthermore, we have found that ISG cells are clonally related to GZMK-expressing CD8+ TILs. Pseudotime trajectory inference suggested a differentiation pathway of ISG > GZMK > Stim-1 cells. However, experimental perturbation revealed that ISG cells are transcriptionally stable and inert to TCR-stimulation. As such, ISG cells may represent a barrier to the differentiation of GZMK cells and subsequent terminally differentiated subsets. Although, numerous unknowns remain and ultimately further experimentation is required to understand the functional implications of this differentiation pathway and these cellular states.

This is not the first report to describe a population of cells enriched with interferon-stimulated genes. Indeed, numerous others have observed similar populations amongst malignant, infectious, and healthy tissues (X. Wang et al. 2022; Quah et al. 2023; Gideon et al. 2022; Cillo et al. 2020). However, the absence of specific cell-surface markers has hindered investigation efforts. Thus far, reports of this population have been limited to mere observation of their appearance. Illustrative of this, Wang and colleagues identified a subset of ISG cells within sequencing data of healthy PBMCs. Despite their efforts, the authors were unable to experimentally isolate this population and thus were limited in the functional analysis that could be performed (X. Wang et al. 2022). Therefore, alternative markers and/or strategies to identify and isolate cells with this cellular state are required. In absence of this, our multimodal sequencing and experimental perturbation approach has provided novel insights into ISG CD8+ TILs.

*Ex vivo* stimulation additionally revealed a cluster of cells that predominantly arose from ISG and GZMK clusters. These clusters had substantial clonal overlap and trajectory inference suggested ISG cells transition through a GZMK phenotype towards the fully activated T-cell state. However, further interrogation of the clonal response to stimulation revealed that ISG cells are transcriptionally inert to TCR-stimulation. The relationship between GZMK and ISG cells is notable as others have demonstrated GZMK expression within solid tumors is associated with improved patient outcomes (Rooney et al. 2015; C. Zheng et al. 2017). Although, the nature of this association is unclear, as GZMK is usually correlated with innate cells and naïve phenotypes. For example, GZMK is more dominantly expressed within immature NK cells. However, GZMK expression within CD8+ T-cells is predominantly observed within central memory and effector memory subsets (Duquette et al. 2023). Thus, supporting the notion that GZMK expression within CD8+ T-cells may correlate with favourable prognosis. Although, it has been observed that GZMK+ CD8+ T-cells are poorly cytotoxic and instead produce IFNγ (Harari et al. 2009; Duquette et al. 2023). Interestingly, others have differential effects of TCR or cytokine stimulation on GZMK expression. Namely, that TCR-stimulation induces the release of GZMK and increase in GZMB expression. Conversely, cytokine-based stimulation drives accumulation of GZMK (Duquette et al. 2023). These findings are consistent with our results which demonstrated TCR-based stimulation drives GZMK cells to down-regulate GZMK and up-regulate GZMB as they differentiate towards a more terminal effector phenotype. Therefore, these data suggest GZMK positivity marks CD8+ T-cells which are not yet terminally differentiated and instead possess a more memory-like phenotype. Additionally, our data suggests ISG cells differentiate into GZMK cells however, they possess relative transcriptional stability. As such, TCR-stimulation is insufficient to drive ISG cells to adopt a GZMK transcriptional phenotype. Thus, ISG cells could function as a barrier within this differentiation trajectory. The functional consequences of this are unknown. Given the above model, accumulation of ISG cells could prevent the development of more terminally differentiated anti-tumoral responses via GZMK intermediaries. However, GZMK+ CD8+ T-cells have been observed within tumor stroma and have been implicated in poor prognosis (Tiberti et al. 2022). Additionally, GZMK CD8+ TILs have been described as a transition state on the trajectory towards exhaustion (C. Zheng et al. 2017; Sun et al. 2022). This is consistent with reports showing IFN-I signalling as a driver of T-cell exhaustion (Chen et al. 2022; Sumida et al. 2022). Therefore, the functional consequences of ISG and GZMK TILs is poorly defined. Further studies are required to better understand the dynamics and function of T cell clusters infiltrating tumor tissues.

## Methods Patient

### Samples

A total of eight patients who had provided informed consent, were included in this study. Samples were obtained from surgical resections of primary HNSCC tumors. All patients presented with oral cavity squamous cell carcinoma and were confirmed to be human papillomavirus (HPV) negative. Fresh HNSCC tumors were collected at the time of resection of the primary tumor and sampled by a pathologist prior to fixation. Fresh tissue was processed to isolate tumor cells and immune cells prior to preservation and storage in liquid nitrogen. The patients enrolled in this study were treatment naïve and characteristics can be found in Figure 1A. Ethical approval for this study was obtained from the Royal Brisbane and Women’s Hospital Human Research Ethics Committee and the QIMR Berghofer Human Research Ethics Committee, HREC/18/QRBW/245.

### Single-cell RNA sequencing

Cells from each patient were cultured as single-cell suspensions and were either stimulated using CD3/CD28 beads or left unstimulated for a duration of 5 hours. Following culture, the cells were sorted using fluorescence-activated cell sorting to isolate live CD45+CD3+CD4-CD8+ cells. Patient samples were sequenced as two unstimulated and two stimulated samples where each sequencing sample represented a pool of 4 patients. As such, approximately 10,000 cells per sample pool were carried forward into the 10x Genomics Single-cell 5’ library pipeline. The libraries were sequenced using a NextSeq 550 (Illumina). The sequencing was performed at QIMR Berghofer Medical Institute.

### scRNAseq pre-processing

Sequencing reads were processed using cellranger (version 3.1.0) and reads were aligned to human reference genome GRCH38-3.0.0. Output from cellranger was processed using Seurat (version 4.3.0). Each sequencing sample was filtered to keep only cells that had a minimum of 200 features and keep features that were detected in a minimum of 3 cells. Subsequently, the two unstimulated samples were merged and the two stimulated samples were merged to give two Seurat objects. These Seurat objects were further filtered to remove cells with greater than 2,500 features or greater than 10% mitochondrial content. Filtering resulted in 5,785 cells with 15,429 features in the unstimulated dataset and 6,042 cells with 15,618 features in the stimulated dataset. Datasets were normalised using LogNormalisation with a scale factor of 10,000. Subsequently, mitochondrial percentage and nCount variables were regressed out using a linear model. Unstimulated and Stimulated datasets were integrated using the Seurat integration pipeline. Unless otherwise stated integration functions/pipeline was executed using default function variables. Integration anchors were calculated using “cca” reduction, “LogNormalize” as a normalization method, and “rann” as the Nearest Neighbour method. Integration resulted in a dataset of 18,295 features across 11,827 cells.

### scRNAseq analysis

#### Dimension reduction and cluster identification

The top 30 PCAs were calculated on the integrated dataset and Nearest-neighbors computed using the top 20 dimensions. Clusters were determined using a cluster resolution of “0.4”. UMAP in figure 1 was generated using top 20 PCA dimensions, the “uwot” algorithm, n.neighbors = 30, and min.dist = 0.3. Following UMAP dimension reduction calculation, clusters were investigated both with manually curated gene signatures and with the use of SingleR (version 2.0.0) to classify cells using data from celldex (version 1.8.0). Two low abundance clusters were removed that were identified as either having high mitochondrial content or a myeloid signature. UMAP projection was recalculated following the removal of these clusters, using the same parameters as previously stated. Therefore, after cluster identification the dataset contained 20,295 features across 11,658 cells with 5,724 cells from the unstimulated treatment condition and 5,934 cells from the stimulated treatment condition. Subsequently, unconventional T-cell clusters were subsetted from the dataset resulting in unconventional T-cell-only and CD8-only datasets. UMAP projections were recalculated for these datasets using the top 20 PCA dimensions, n.neighbors = 50, and a min.dist of 0.1 for CD8-only dataset or 0.5 for the unconventional T-cell-only dataset. The unconventional T-cell-only dataset consisted of 20,295 features across 970 cells. The CD8-only dataset consisted of 20,295 features across 10,688 cells, 5,165 of which originated from the unstimulated treatment condition and 5,523 from the stimulated treatment condition.

#### Differential gene expression

Calculations to determine differentially expressed genes between clusters or conditions was performed using wilcox test implemented via the standard Seurat analysis pipeline. Analysis was performed using the RNA data slot of the Seurat object.

#### Differentially expressed transcription factors

To determine the differential expression of transcription factors, the list of differentially expressed genes was cross-referenced with a curated database of RNA polymerase II regulated transcription factors (TFcheckpoint; http://www.tfcheckpoint.org).

#### Gene ontology analysis

Briefly, differentially expressed genes for the ISG cluster were identified using Seurat’s FindMarkers() function. Genes identified as significantly (adjusted p.value < 0.5) up- or down-regulated were then passed to the enrichR package (version 3.1) to identify enriched terms using the GO_Biological_Process_2021 database. The top 10 enriched terms were then visualised using SCpubr (version 1.1.1).

#### Signature scoring

Signature score calculated using UCell (version 2.2.0) with signatures for type I and II IFN obtained from (Azizi et al. 2018).

#### Cell-cell communication analysis

Cell-cell communication was performed using the R package “liana” (version 0.1.12). In brief, cell-cell communication networks were calculated using the following methods “natmi”, “connectome”, “logfc”, “sca”, and “cellphonedb”. The scores from these methods were subsequently aggregated and only interactions concordant between methods were kept. This analysis followed the recommended analytical pipeline for the “liana” package.

#### Trajectory Inference

Trajectory inference was performed using the dynverse (Cannoodt and Saelens 2023) collection of packages. Analysis was performed using standard pipeline with default parameters and without supplying any priors for both slingshot and Scorpius trajectory inference algorithms.

### scRNAseq visualisation

#### Imputation

Imputation of gene expression was performed and used in certain visualisations where indicated. Imputed values were not used for any downstream analysis and were exclusively used in indicated visualisations. Imputation was performed using the “RunALRA” function in Seurat and increased the percentage of non-zero values in the dataset from 29.63% to 38.95%.

#### Density based UMAP visualisation

The Nebulosa package (version 1.8.0) together with scCustomize package (version 1.1.1) were used to visualise gene expression on UMAP projections and expression density.

#### Color scheme

Where possible the uniform, colorblind-friendly batlow (Crameri, Shephard, and Heron 2020) color pallet was used for data visualisation. The color palette was accessed using the scico package (version 1.3.1).

### Single-cell TCR sequencing analysis

#### Pre-processing

Single-cell TCR sequencing data were aligned using cellranger pipeline (version 3.1.0) to the human VDJ reference (vdj_GRCh38_alts_ensembl-3.1.0-3.1.0). TCR data was subsequently processed using scRepertoire (version 1.8.0). TCR data was filtered such that if cells had multiple alpha or beta chains identified, only the top expressing chain was retained. Additionally, unless otherwise stated, clone identity was defined by the CDR3 amino acid sequence.

#### Clone size definitions

Abundance of clones was calculated per stimulation condition and binned according to the following definitions. Single (x = 1), small (1 < x < =5), medium (5 < x <=10), large (10 < x <=20) and hyperexpanded (20 < x <= 150). Where x = number of cells with exact CDR3 amino acid sequence. Size cut-offs were determined empirically using summary statistics of clone abundances across the dataset.

### External datasets

#### uTILity

The pan-cancer “uTILity” dataset was acquired from (Nicholas Borcherding 2022) circa 13.10.2022. The dataset was filtered to retain only cells identified as CD8 T-cells and only Tumor and Normal tissue types were retained. Subsetted dataset was normalized and reintegrated using the harmony package (version 0.1.1) to remove “Cohort” effect. UMAP coordinates and clusters were recalculated following harmonization, using the standard Seurat analysis pipeline.

#### HNSCC

For validation of ISG gene signature and cell-cell communication analysis, the HNSCC TILs dataset published in (Cillo et al. 2020) was used. Processed data was downloaded from (GSE139324)[https://www.ncbi.nlm.nih.gov/geo/query/acc.cgi?acc=GSE139324]. Metadata for this dataset was obtained through contact with the lead author/s.

#### COMBAT dataset

The Covid-19 and Infleunza scRNAseq dataset was downloaded from https://zenodo.org/records/6120249 (COvid-19 Multi-omics Blood ATlas (COMBAT) Consortium 2022)

## Figure preparation

Figures were arranged and formatted using Adobe Illustrator (version 27.5) and/or GraphPad Prism (version 9).

**Table 1:**
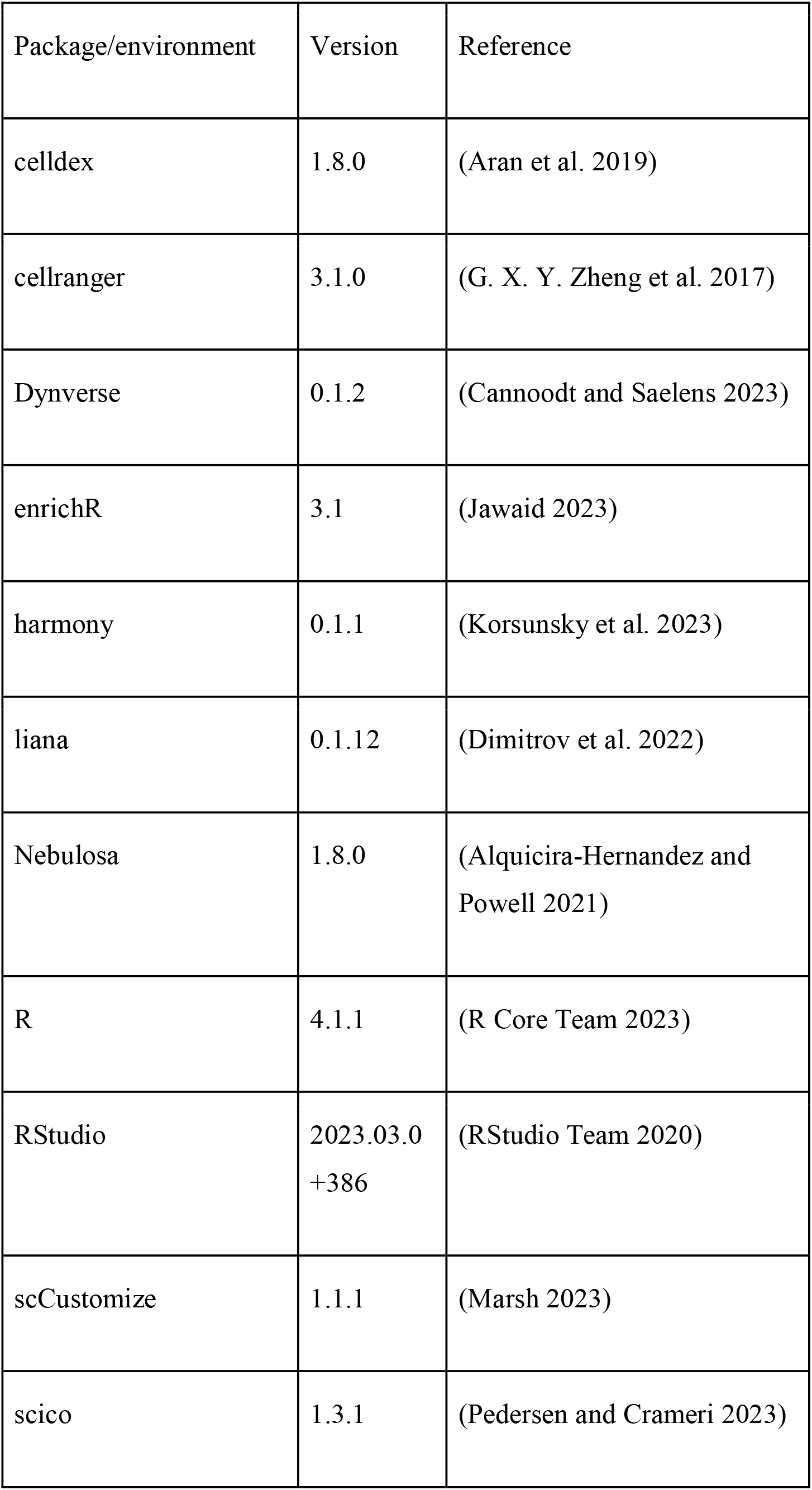
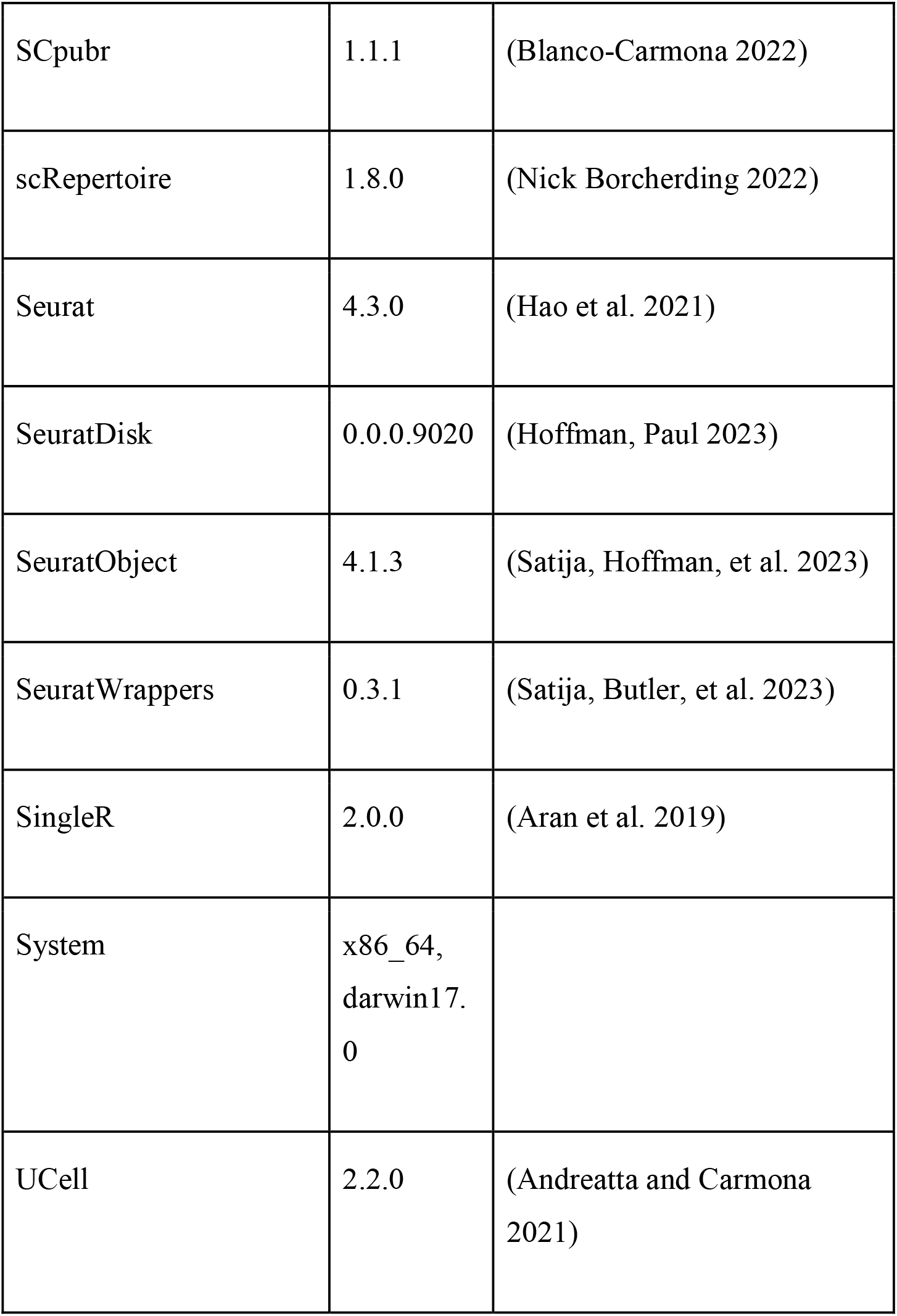
Analysis packages used.

Table depicting the analysis packages and the software environments used within this manuscript

## Availability of data and materials

The single-cell RNA/TCR sequencing dataset generated will be made available upon reasonable request and approval of HREC. All code used to generate figures can be found under the relevant repository at https://github.com/BaldLab. All other data generated are available upon request.

## Authors contributions

Conceptualization – D.C, M.B and T.B

Methodology – D.C, M.B,

Software – D.C, T.K, L.M.S, D.P, N.B

Formal Analysis – D.C, T.K, L.M.S, D.P, N.B

Investigation – D.C, N.S, M.B, T.B

Resources – N.B, M.H, T.B, K.T, S.P, Ma.Ba, J.M, B.H

Data curation – D.C, L.T.K, N.B

Writing (Original Draft) – D.C, M.B, T.B

Writing (Review & Editing) – all authors Visualization – D.C, N.B

Supervision – D.C, M.B and T.B

Funding acquisition – T.B

Project administration – T.B

## Acknowledgements

Firstly, we wish to extend our appreciation for the patients whom provided their samples, without which, this study would not have been possible. Additionally, we express our gratitude to the flow cytometry and next generation sequencing facilities of QIMR Berghofer Medical Research Institute. We thank aimed analytics GmbH for bioinformatics support. We thank Christian Engwerda from QIMR Berghofer Medical Research Institute for advice and support on this manuscript.

