## Supplementary figures and images for "Type I interferon drives a cellular state inert to TCR-stimulation and could impede effective T-cell differentiation in cancer"

### Supplemental Figure 1

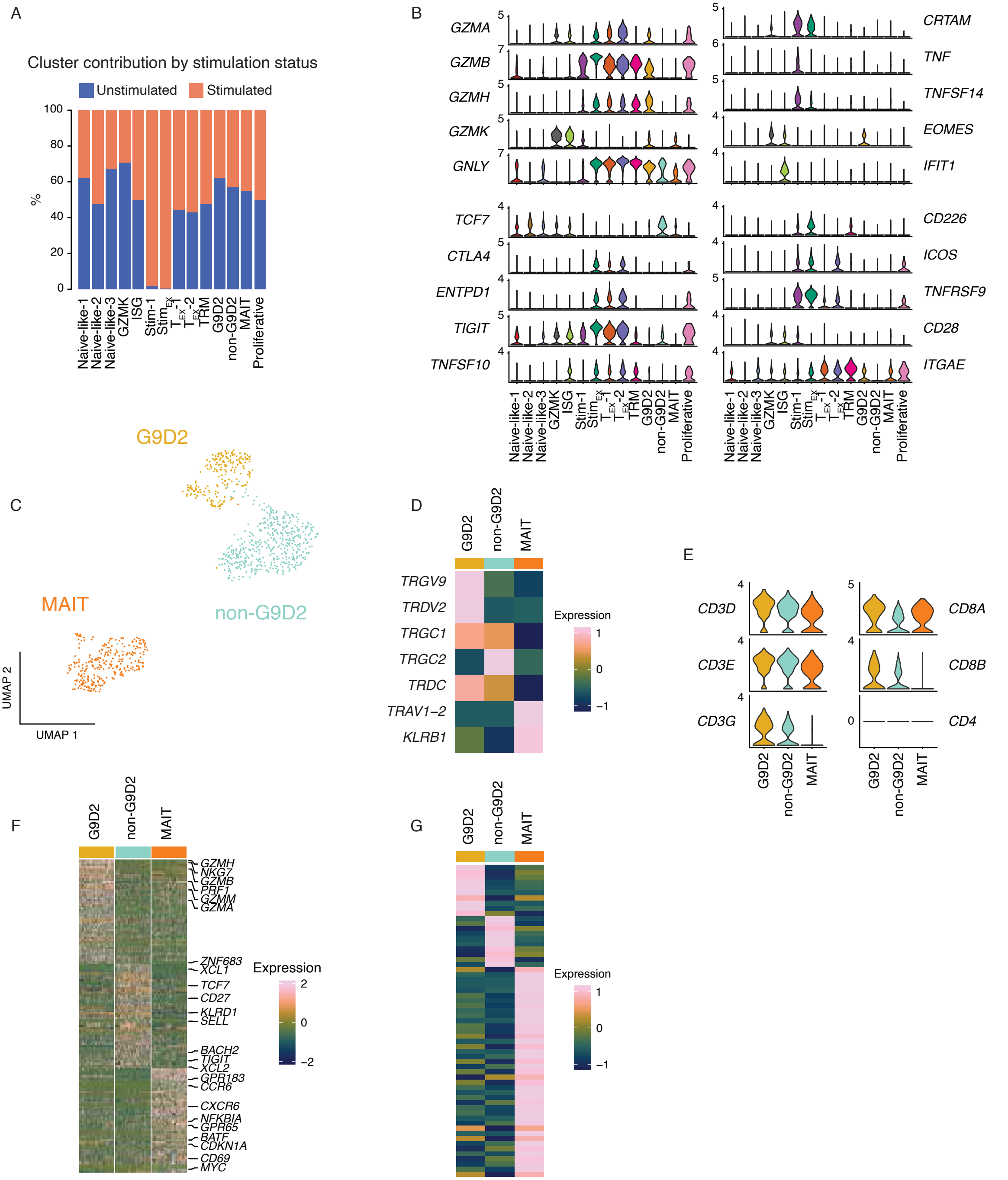

### Supplemental Figure 2

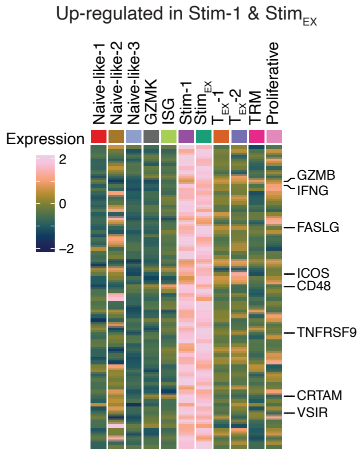

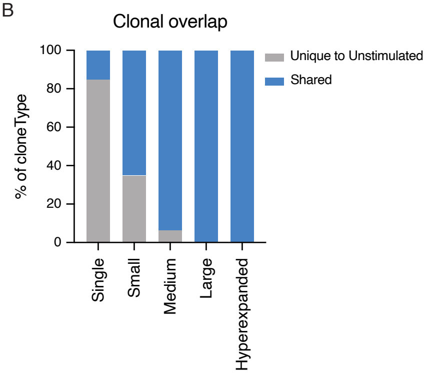

### Supplemental Figure 3

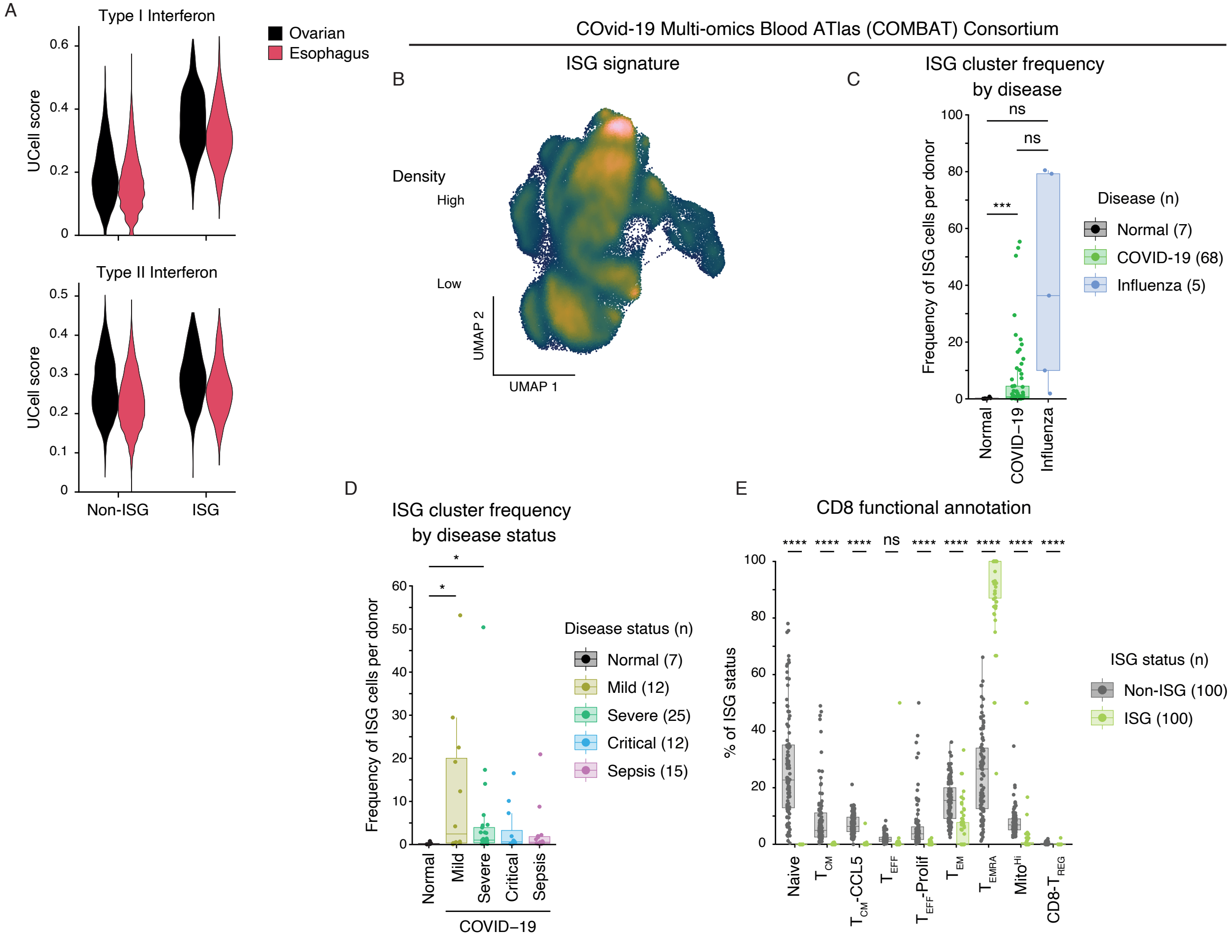
